# Uncovering Minimal Pathways in Melanoma Initiation

**DOI:** 10.1101/2023.12.08.570336

**Authors:** Hui Xiao, Jessica Shiu, Chi-Fen Chen, Jie Wu, Peijie Zhou, Sahil S. Telang, Rolando Ruiz-Vega, Qing Nie, Arthur D. Lander, Anand K. Ganesan

## Abstract

Cutaneous melanomas are clinically and histologically heterogeneous. Most display activating mutations in *Braf* or *Nras* and complete loss of function of one or more tumor suppressor genes. Mouse models that replicate such mutations produce fast-growing, pigmented tumors. However, mice that combine *Braf* activation with only heterozygous loss of *Pten* also produce tumors and, as we show here, in an Albino background this occurs even with *Braf* activation alone. Such tumors arise rarely, grow slowly, and express low levels of pigmentation genes. The timing of their appearance was consistent with a single step stochastic event, but no evidence could be found that it required *de novo* mutation, suggesting instead the involvement of an epigenetic transition. Single-cell transcriptomic analysis revealed such tumors to be heterogeneous, including a minor cell type we term LNM (<u>L</u>ow-pigment, <u>N</u>eural- and extracellular <u>M</u>atrix-signature) that displays gene expression resembling “neural crest”-like cell subsets detected in the fast-growing tumors of more heavily-mutated mice, as well as in human biopsy and xenograft samples. We provide evidence that LNM cells pre-exist in normal skin, are expanded by *Braf* activation, can transition into malignant cells, and persist with malignant cells through multiple rounds of transplantation. We discuss the possibility that LNM cells not only serve as a pre-malignant state in the production of some melanomas, but also as an important intermediate in the development of drug resistance.

## INTRODUCTION

Melanomas typically exhibit DNA alterations involving multiple oncogenes and tumor suppressor genes^1–5^. However, because melanocytes—the cell of origin of melanoma—usually possess a high background of mutation, it has been difficult to identify a specific sequence of events that inexorably leads to melanoma^6^. Whereas activating *BRAF* mutations are the most commonly observed driver mutations in melanomas, occurring in 60% of cases ^7^, the same mutations are also found in nearly all growth-arrested melanocytic nevi ^8^. This result, together with the observation that expressing activated *Braf* in mouse melanocytes primarily generates nevi, has led to the view that MAP kinase pathway activation (through mutation of *Braf* or, alternatively, *Nras*) is necessary to initiate melanoma but generally not sufficient.

Consistent with this view, when activated *Braf* (or *Nras*) in mouse melanocytes is combined with complete loss of function mutations of tumor suppressor genes (*Pten*, *p53*, *Ink4a*, *Cdkn2a*), melanomas arise rapidly ^9–14^. It has been hypothesized that, without the help of these additional mutations, activation of *Braf* or *Nras* leads to oncogene induced senescence (OIS) and cell cycle arrest, explaining why only nevi are formed^15–19^. Recent studies cast doubt on this hypothesis, however, and show both that *Braf* activation does not induce melanocyte senescence *in vivo*^20^, and that nevus cells remain competent to proliferate^21^. At present, the mechanism behind growth arrest following *Braf* activation remains unknown but may involve feedback loops normally involved in melanocyte homeostasis^20^.

When activated *Braf* is expressed in the melanocytes of mice, in some cases (depending on the expression construct and genetic background^10,13^), rare melanomas do arise, amongst a background of abundant nevi. While it seems plausible that the spontaneous acquisition of additional mutations explains such malignant transformation, that hypothesis has not been tested. Indeed, such melanomas are far less studied than either human melanomas, or mouse melanomas produced when multiple mutations (typically three or more) are introduced simultaneously^22^. While mouse melanomas that combine many mutations can be expected to model advanced human disease, by the same token, they are not necessarily a good model for elucidating the stepwise progression that occurs between normal melanocyte and melanoma.

Recent studies suggest that such progression is likely to involve not only the accumulation of mutations but also non-genetic transitions among cell states. Melanomas display a high degree of intratumoral heterogeneity, both in cellular behaviors and epigenetic modification ^23,24^, and stochastic, non-genetic switching between melanoma cell states has been directly observed *in vitro* ^25–29^ and proposed to occur *in vivo*. In some cases, it has been argued that specific states serve as necessary waystations on the path to therapeutic resistance^30,31^. Little is known, however, about what drives transitions between states, nor the extent to which mutational versus non-genetic processes influence them.

With the advent of methods for assessing transcriptomes at the single cell level, several groups have attempted to define the landscape of cellular heterogeneity in melanoma. Based on analyses of human melanomas, mouse models, and cell lines, gene expression signatures have been proposed to define cell states that have been variously categorized as pigmented, invasive, proliferative, neural crest-like, mesenchymal, intermediate, and *BRAF* inhibitor resistant^28,31–34^. Signatures derived in different studies are often divergent, and it is unclear whether this reflects real biological differences or variations in data quality or informatic approaches. A complicating factor is that most data come from the analysis of melanomas bearing different combinations of driver mutations (including in some cases ones that were not fully characterized) in a variety of genetic backgrounds.

Here we sought to clarify the cellular complexity of melanoma, particularly during early stages of tumor development, by focusing on mouse models of “minimal” induction, i.e., models in which melanocytes are engineered to express only activated *Braf*, or activated *Braf* plus a single null allele of the tumor suppressor *Pten*. On a genetic background that allowed for rare melanoma emergence even when only *Braf* was mutated, the kinetics of tumor appearance were consistent with a requirement for a single stochastic event. Yet, surprisingly, we could find no evidence that this event involved DNA mutation. Analysis by single cell RNA sequencing of tumors, as well as adjacent and normal skin, revealed that the major cell types in minimally-mutated melanomas expressed relatively low levels of pigmentation genes, and the resulting tumors were indeed hypomelanotic. Through direct comparison of gene expression signatures, it was possible to identify major cell states in such tumors that correspond reasonably well to minor populations observed in more heavily mutated melanomas, including a state specifically thought to play a role in the development of therapy resistance^31^. Intriguingly, one of the states we observed was also detected in tumor-free skin, suggesting it is not obligatorily malignant when it is present alone. Using new informatic tools we predicted that this cell may act as a direct source of more abundant types of tumor cells. We discuss the possible relationship of cell states we identify here to events in human melanoma initiation and progression.

## RESULTS

### *Braf* activating mutations induce rare melanomas in both *Pten*-heterozygous and wild-type albino mice

We used a *Tyrosinase::CreERT2; Braf^CA-fl/+^* construct^35^ in transgenic mice to specifically activate *Braf* in melanocytes by topically applying tamoxifen at postnatal days 2-4 (Fig 1A). On a C57/Bl6 background, such mice developed abundant nevi in a matter of weeks, but no melanomas. In contrast, when a single floxed allele of *Pten* was introduced into this background, creating a heterozygous *Pten* mutation in melanocytes, sporadic tumors arose, an average of three per animal (Fig 1A, S1A-B). Both gross (Fig 1A, S1A) and histologic (Fig 1C) analysis of these tumors strongly suggest they did not arise by loss of heterozygosity of the wild type *Pten* allele. This is because melanomas generated by activating *Braf* together with loss of both *Pten* alleles (*Pten^Δ/Δ^*) are strongly pigmented^13,36^ (Fig 1C, right lower image), whereas those arising on a *Pten^Δ/+^*background have a very different appearance, displaying very little pigment either visually (Fig 1A, S1A) or histologically (Fig 1C).

**Figure 1.**
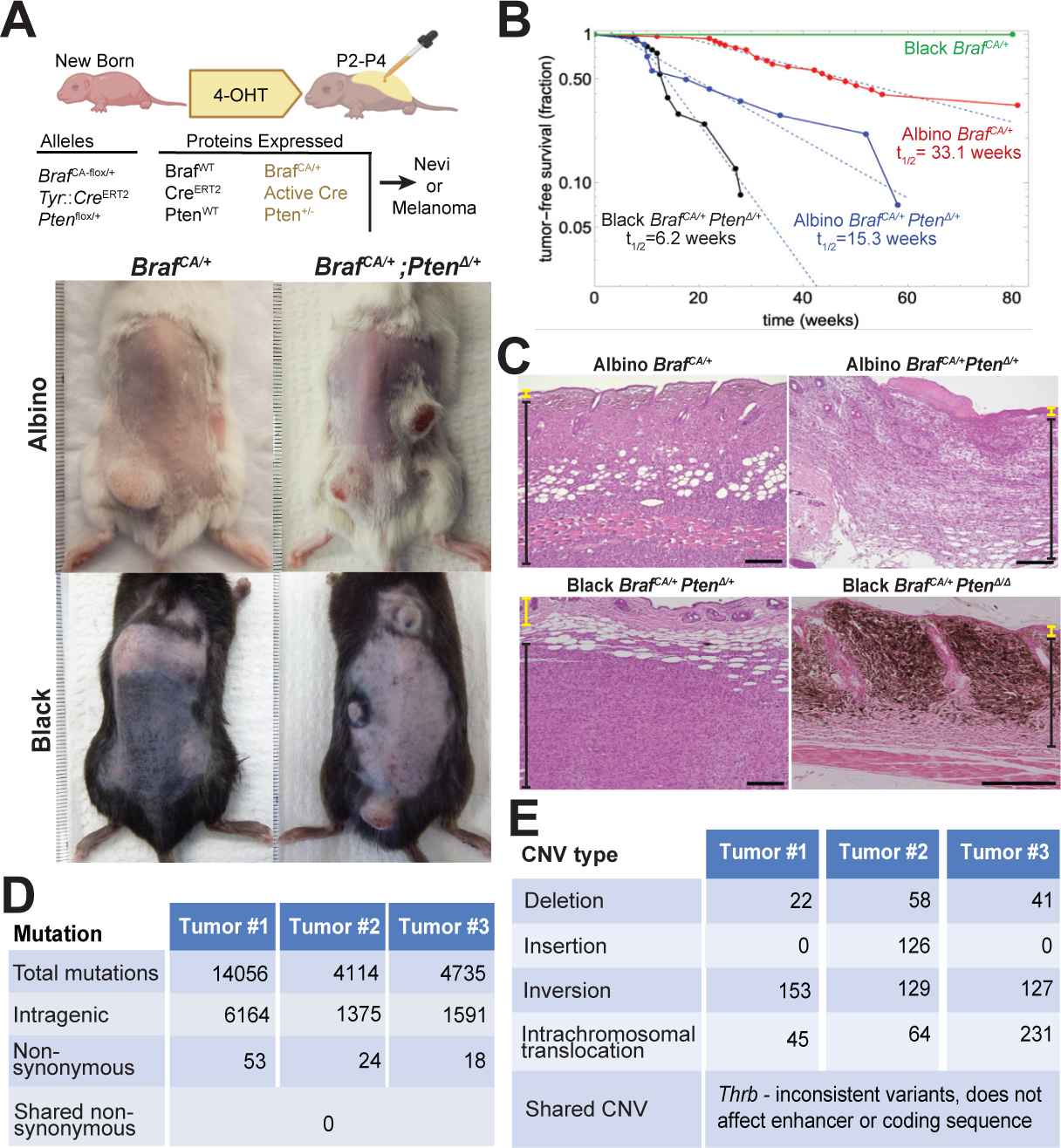
The *Braf*^CA/+^ mutation can induce melanomas in mice independently of additional mutations. **A.** Images and **B.** tumor-free survival curves for *Braf*^CA/+^ and *Braf*^CA/+^; *Pten*^Δ/+^ mice in different coat-color backgrounds. Albino *Braf*^CA/+^ mice develop tumors not observed in congenic Black *Braf*^CA/+^ mice. **C.** H&E staining of tumors. Yellow and black brackets on the side denote tumor-free and tumor-containing regions, respectively. Note that different tumor genotypes differ markedly in pigmentation. *Pten*^Δ/Δ^ tumors are highly pigmented, especially superficially, whereas *Pten*^Δ/+^ tumors are more infiltrative and largely lack pigment. Scale bar: 200um. **D.** Whole-genome sequencing (WGS) was performed on three Albino *Braf*^CA/+^ tumors (80X coverage). Observed mutations (intragenic and non-synonymous), including single nucleotide variant (SNV), insertion, deletion, and multi-nucleotide variant (MNV) are reported. No non-synonymous mutations were shared among the tumors. **E.** The nearest gene bodies to detected copy number variants in albino *Braf*^CA/+^ tumors were determined. Only one gene, *Thrb*, has CNVs close to the gene body in all three tumors. The locations of these CNVs did not suggest functional significance and did not overlap (see Fig S1C).

We also examined the outcome of expressing *Braf^CA/+^*in the melanocytes of albino mice (congenic with C57Bl/6). Epidemiological studies indicate human melanoma risk is higher in light-skinned than dark-skinned individuals and factors other than tumor mutation rate are known to contribute to this difference^37^. We found that, on an albino background, *Braf^CA/+^* alone produced melanoma in about 2/3 of animals, with no animals displaying more than one tumor (Fig 1A, S1A-B). Combining *Braf^CA/+^*with *Pten* heterozygosity in albino mice increased the numbers of tumors per animal to a level similar to that seen in black *Braf^CA/+^ Pten^Δ/+^* mice (Fig S1A-B).

The rarity of the tumors arising when *Braf* is activated either alone or together with a single loss of function allele of *Pten* suggests that at least one additional event is required to produce melanoma. The distribution of times at which tumors become detectable in such mice strongly suggest that the event is statistically random, i.e. occurs with a constant, rare probability. As shown in Figure 1B, plots of tumor-free survival over time are fit by single declining exponential functions, consistent with “single-hit kinetics”, i.e. the expectation of a process with a constant probability per unit of time. Because of the need for tumors to exceed a threshold size before being detected, we cannot rule out the possibility of *more* than one random event being required for tumor initiation, but based on the application of the single-hit model, we may extract from the final slopes of logarithmic plots a “half-time” (t_1/2_) for tumor initiation (the expected time for tumors to arise in 50% of animals) equal to ∼33 weeks in albino *Braf^CA/+^* mice, ∼15 weeks in albino *Braf^CA/+^ Pten^Δ/+^* mice, and ∼6 weeks in black *Braf^CA/+^ Pten^Δ/+^* mice. It is important to note that, on such plots, the rate of tumor growth is reflected only in the initial lag-time and not the eventual slope, which is a measure of when initiating events occur. Thus, the results suggest that pigmentation status and *Pten* gene dosage both influence the probability of melanoma initiation, consistent with observations in humans, where skin color modifies melanoma risk by mechanisms independent of mutation.^37^

To investigate whether the random, initiating event required to produce albino *Braf^CA/+^* tumors is a *de novo* DNA mutation, we performed whole genome sequencing (WGS) on three such tumors (Fig 1D-E). Tumors were dissected from the overlying skin, and DNA was prepared along with control DNA from spleen and skin of the same animals. Sequencing was performed to an average of 80X coverage (within mappable regions, >99% of nucleotides were covered at least 40 fold). At that depth, even heterozygous variants that were present in 20% of cells would be detected with a minimum of 98% accuracy. A Bayesian somatic genotyping model (Mutect2) was used to identify tumor-specific somatic variants as those present in tumor samples but absent from the spleen of the same animal. Based on the levels of unrecombined and recombined *Braf^CA/+^* alleles detected in each sample, we determined that the fraction of DNA that was tumor-derived in each sample was 19%, 55%, and 36%, for tumors labeled 1, 2, and 3, respectively, in Figure 1D-E. Thus, any mutation that was required for malignant transformation would very likely have been detected in each sample.

Overall, we observed between 4,000 and 14,100 mutations per sample, including single nucleotide variants (SNVs), multiple nucleotide variants (MNVs), insertions, and deletions, many of which were present at too low a frequency to be causal in tumorigenesis (Fig 1D, Tables S1-3). Very few of these variants were intragenic, and only a handful of those produced nonsynonymous changes. Of these, no gene was mutated across all three tumors. In addition, we specifically checked, and did not find, mutations in sequences upstream of *Tert* that correspond to sites of promoter mutations in human melanoma^38^.

We also used paired-end reads to identify breakpoints and infer copy number variations (CNVs) and structural variations (SVs). We used the *BreakDancer* pipeline to identify CNVs that were unique to the tumor and not observed in the spleen or skin of the same animal with a confidence score cutoff of p<.001 and a false discovery rate of 0.05 (a lenient cutoff). We grouped CNVs by the gene bodies to which they were closest. Only one gene, *Thrb,* had CNVs near the gene body in all three tumors sequenced (Fig 1E, Tables S4-6), and the three observed mutations were all small, intronic, and did not overlap, and thus seemed unlikely to have an impact on transcript expression (Fig S1C). We also did not observe expression of *Thrb* in the neural crest derived clusters in our single cell gene expression dataset (Fig S2B), suggesting it would be unlikely for a mutation in this gene to contribute to tumor initiation. Of note, we did not observe CNVs in genes that were identified to have copy number variations in human *Braf* mutant tumors including but not limited to the known tumor suppressor genes *Pten, Cdkn2a, Tert, Nf1, Tp53, Rac1* or *Cdk4*^4^.

#### Identifying Melanocytic Cell States

Taken together, these data suggest that a de novo mutation is unlikely to be the initiating event for tumors in albino *Braf^CA/+^* animals, an observation that raises the possibility that a non-genetic transition—e.g. epigenetic cell state switching or collective cell transition—drives melanoma initiation. Interestingly, non-genetic, stochastic switching between cell types has been described among melanoma cells *in vitro*^29^. If such switching plays a role in tumor initiation in the *Braf^CA/+^* and *Braf^CA/+^ Pten^Δ/+^* mouse models, we reasoned that evidence of it might be found by analyzing the cell types or states within tumors. For example, we might observe a cell state in melanoma tumors that was also found in skin in which tumors had not yet arisen.

To explore this possibility, we performed single cell RNA sequencing on a total of 47 skin and tumor samples from 36 animals: 15 wildtype skin samples, 10 black *Braf^CA/+^* skin samples (which contain only nevi), 3 albino *Braf^CA/+^* tumors, 4 albino *Braf^CA/+^ Pten^Δ/+^*tumors, 4 black *Braf^CA/+^ Pten^Δ/+^* tumors, and 11 sample-matched tumor-adjacent-skin samples collected alongside each of the 11 tumor samples (Fig 2A). After combining samples using the merge function of Seurat, performing quality control, and normalizing using scTransform, a total of 345,427 cells were identified. Using unsupervised clustering, based on highly variable genes and marker genes known to identify cell types, we identified 18 cell types, including melanocytes (*Dct*+, *Pmel*+, *Mitf*+, and *Mlana*+), Schwann-cells (*Mpz*+, *Dhh*+), fibroblasts (*Pdgfra*+, *Col3a1*+, *Sparc*+) and macrophages (*Adgre1+, Cd74+, Ctss+*) (Fig 2B, Fig S2A, Table S7). Specifically within tumor samples, an abundant cell type expressed markers commonly seen in melanoma, including the markers *Plp1, Gpm6b, Postn, Mcam, S100b*^39^ (Fig 2B, 2D).

**Figure 2.**
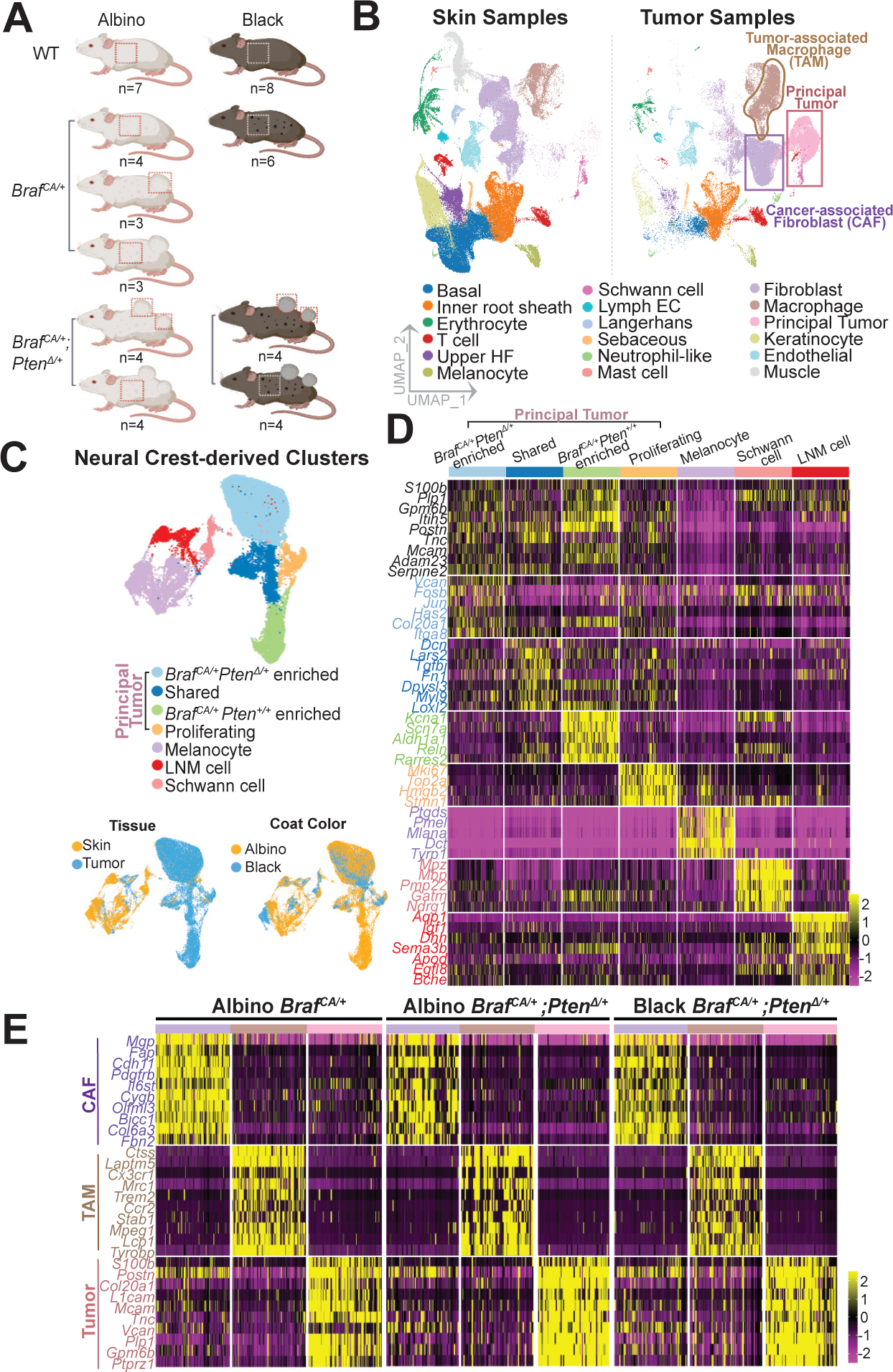
Single-cell transcriptomics identifies melanocyte/neural crest-derived, macrophage, and fibroblast populations in normal skin, *Braf*^CA/+^ and *Braf*^CA/+^; *Pten*^Δ/+^ tumors. **A.** Coat colors, genotypes and numbers of mice subjected to single-cell RNA sequencing. **B.** 345,427 cells from 36 mice (47 normal, nevus-containing skin, and melanoma samples) from the genotypes in panel A were subjected to scRNAseq, jointly clustered, and projected onto a common UMAP. ScRNA-seq identifies populations enriched in tumors, highlighted in the right panel. HF: Hair follicle. EC: Endothelial cell. **C.** Neural crest-derived clusters (identified by the expression of *S100b*, *Dct*, *Mpz* [as well as a Cre-reporter gene in selected cases]) were further subsetted (35,527 cells in total) and populations specific to tissue and coat color were characterized (bottom panel). **D.** Differentially expressed genes associated with neural crest-derived clusters defined in panel C. Known melanoma markers are labeled in black; colored bars and other text colors correspond to labeling in C. **E.** Gene expression profiles of tumor-enriched fibroblasts (purple), macrophages (brown) and tumor cells (pink) compared among the tumor genotypes.

We then subsetted the cells in all samples that were judged to have a potential lineage relationship to melanocytes. These included melanocytes themselves, Schwann cells (neural crest-derived cells thought to share a common precursor with melanocytes ^40,41^), and any cell types in which we could document recombination driven by the *Tyr*-*Cre^ERT^*^2^ transgene (in a subset of samples, the mTmG reporter system ^42^ was used so that expression of GFP marked cells in which Cre-mediated recombination had occurred); the latter included the abundant cell type in tumor samples that expressed melanoma markers.

These 35,527 cells, which we refer to as “neural crest derived”, were re-normalized using ScTransform and subclustered (Fig 2C). We identified within this group melanocytes, which produce a high level of pigmentation genes (*Ptgds*, *Pmel*, *Mlana*) consistent with the highly pigmented nature of nevus melanocytes ^20^; a Schwann cell population (marked by *Mpz*); and several tumor-enriched cell populations that clustered separately from melanocytes, and expressed melanoma markers (*Plp1, Gpm6b, Postn, Mcam*^39^) (Fig 2C-D). A distinct population of cells was observed primarily, but not exclusively, in tumor samples and could be distinguished by weak but detectable expression of pigmentation genes (*Mlana, Tryp1*), expression of multiple “neural” genes (*Bche, Sema3b*), and expression of genes encoding structural components of the extracellular matrix. We termed these LNM (<u>L</u>ightly pigmented, <u>N</u>eural, extracellular <u>M</u>atrix) cells. Other markers of LNM cells included *Aqp1, Dhh,* and *Igf1* (Fig 2D, Fig S2B). Although LNM cells were relatively abundant in tumor samples, they were also observed in normal skin, and their numbers increased substantially in the tumor-free skin of *Braf^CA/+^* mice (Fig S2C). We calculated the Euclidean distance in principal component space between LNM cells, the principal tumor cluster, Schwann cells and melanocytes and observed that the LNM cells were closer to principal tumor cells than to the Schwann cells or melanocytes (Fig S2D, Table S8). Immunohistochemical staining for *Aqp1* indicated that LNM cells were sparse in normal skin, located in or around nevi in black *Braf^CA/+^* skin, and were absent in the upper dermis but filled the mid and lower dermis of the *Braf^CA/+^* tumors (Fig S2E).

Principal tumor cells, defined by the expression of known melanoma marker genes *Postn*+, *Mcam*+, *Plp1*+, *Gpm6b*+, ^39^ were observed in tumors from all three genotypes (Fig 2E). These cells could be subclustered into four distinct subsets: a cluster with a gene expression signature that was shared amongst tumor types (shared), a cluster with a proliferative gene expression signature that was shared amongst tumor types (proliferating), a cluster with a gene expression signature unique to *Pten^Δ/+^* tumors, and a cluster with a gene expression signature unique to *Pten^+/+^* tumors (all of the latter were also albino) (Fig 2C-D, S2B). The clusters observed in all genotypes also expressed extracellular matrix related genes (*Vcan, Col20a1*) and exhibited low level expression of pigmentation genes (*Mitf*, *Dct*, *Pmel*)^43^ (Fig 2D,2E) consistent with the observation that pigment in these tumors was not readily apparent visually or histologically even in black mice (Fig 1A, C, S1A). The proliferating cluster was marked by the expression of proliferation markers of *Top2a*^44^, *Mki67*^44^, *Hmgb2*^45^ (Fig 2C-D, S2B). The *Pten^Δ/+^*-specific signature was distinguished by expression of genes associated with cell stress (*Fos, Jun*)^46^. The albino *Pten^+/+^-*specific cluster was marked by expression of genes that others have noted to mark neural crest stem cells (NCSCs) (*Aldh1a1, Ngfr, Reln*) ^28,47^ (Fig 2D, Fig S2B).

We also identified and subclustered fibroblastic and immune cells in both skin and tumor samples. Cancer-associated fibroblasts (CAFs) and tumor-associated macrophages (TAMs) were specific to tumor tissue (Fig 2B); their gene expression profiles were consistent across tumor genotype (*Braf^CA/+^ Pten^+/+^* vs. *Braf^CA/+^ Pten^Δ/+^*) (Fig 2E). CAFs expressed markers that had been previously identified in other cancer types (*Mgp*^48^, *Igfbp7*^49^, *Sparcl1* ^50^, *Col4a1*^51^, *SerpinE2*^52^) (Fig S3A). TAMs shared the expression of some macrophage specific markers with tissue macrophages, but failed to express other markers characteristic of tissue macrophages (*Cxcl2*^53^ and *Retnla*^54^ (Fig S3B)).

### LNM Cells Persist after Tumor Transplantation

Single cell RNA sequencing identified LNM cells as a population present at low levels in normal skin, enriched in *Braf^CA/+^*skin (Fig S2C) and highly enriched in tumors (Fig S2E). To assess whether LNM cells are an integral part of the tumor microenvironment, we subjected tumors to serial transplantation, using an approach developed for propagating patient derived xenografts^55^. Black *Braf^CA/+^ Pten^Δ/+^*tumors were readily transplantable, with the highest rate of success occurring when immune deficient (NSG) hosts were used. Interestingly, the time delay to initiate the growth of melanoma tumors shortened after rounds of serial transplantation (Fig S4A), consistent with enrichment for tumor generating populations. Single cell transcriptomics was used to quantify numbers of cells of different types after different rounds of transplantation (Fig 3A). After merging single cell data from transplants and the primary tumors from which they derived, and clustering by gene expression, we identified four neural crest-derived clusters: a principal tumor cluster, a melanocyte cluster, and two LNM clusters (Fig 3B-C, S4B). Interestingly, the number of LNM cells increased after successive rounds of transplantation (Fig 3C), from 11% to 74% of neural crest derived cells. We also observed a shift in the gene expression signature of some LNM cells after multiple rounds of transplantation, which led to LNM cells being divided into two clusters. This might either have been the result of selection or a response to growth in an immune-deficient environment. Nonetheless, when we calculated the Euclidean distance between all neural crest-derived cell types we observed that both of the LNM populations were approximately equidistant between melanocytes and tumor cells (Fig S4C, Table S9).

**Figure 3.**
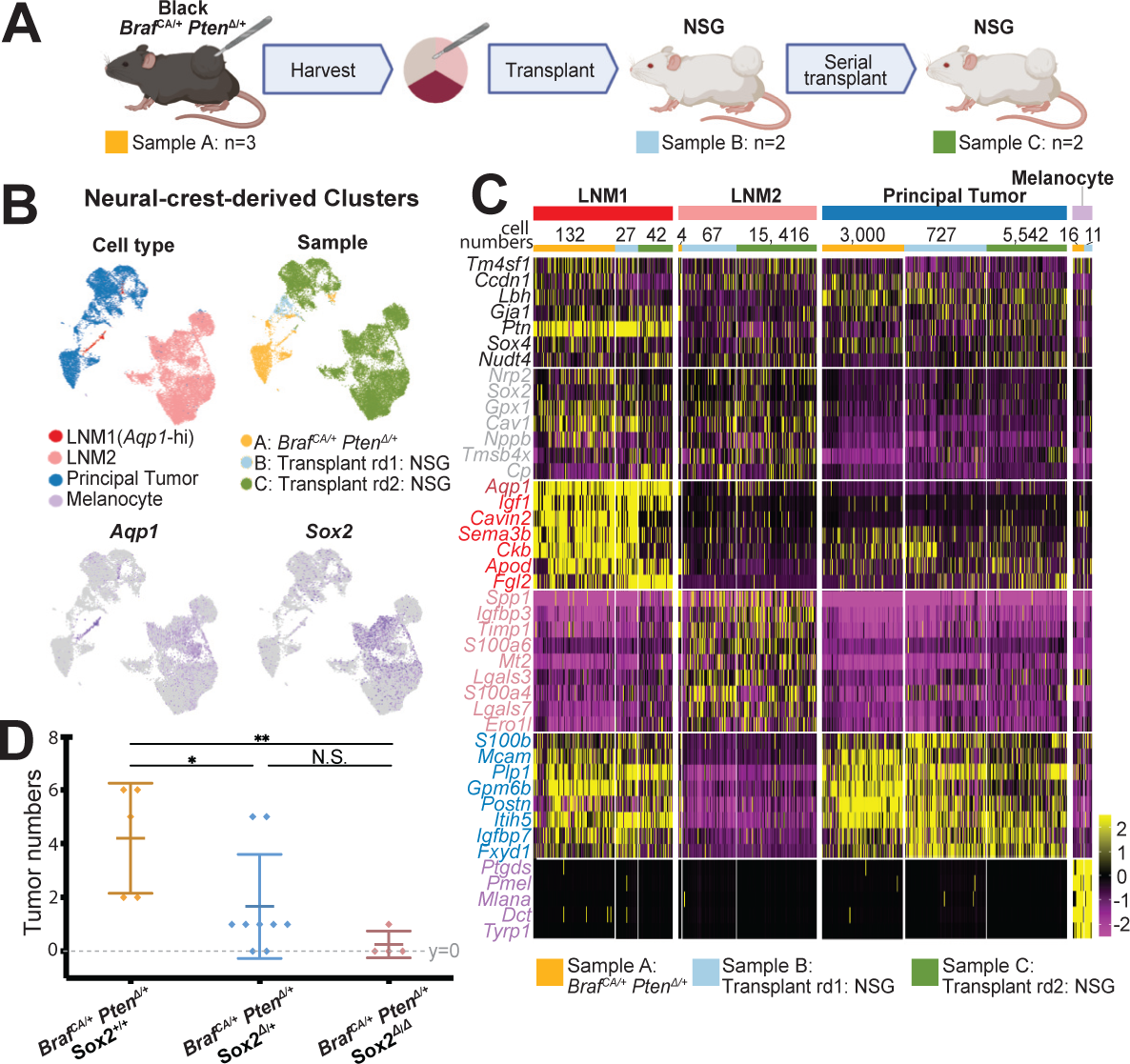
LNM cells persist after transplantation. **A.** Tumors derived from *Braf*^CA/+^; *Pten*^Δ/+^ were serially passaged by transplantation in to NSG (immune deficient) mice. **B.** Single-cell RNA sequencing and subclustering was used to identify neural crest-derived clusters (24,984 cells), among which both principal tumor and LNM clusters were identified. rd = Round of transplantation. **C.** Gene expression profiles of main cell types in panel B. Gene signatures shared among LNM1, LNM2, and Principal Tumor are labeled in black, and a signature shared by LNM1 and LNM2 are labeled in grey. **D.** Conditional *Sox2* deletion in melanocytic/neural crest derived cells inhibits tumor development. Tumor incidence in mice of the indicated genotypes is shown.

### A role for *Sox2* in melanoma initiation

We noted that LNM cells were specifically enriched in the expression of the stemness-associated transcription factor *Sox2* (Fig 3C). *Sox2* is also detected in human melanomas^56^ and has been described as enriched in melanoma-initiating cells^57^, but studies in mouse models have so far failed to support a role for it in melanoma formation. Specifically, in mice bearing either combined *Braf^CA/+^ Pten^Δ/Δ^* ^58^ or *Nras^Q61K/+^ Ink4a ^Δ/Δ^* ^59^ mutations, loss of *Sox2* did not prevent tumor formation. When we crossed conditional alleles of *Sox2* (*Sox2^fl^*) into the *Braf^CA/+^ Pten^Δ/+^* background, however, we observed a different result. When *Braf^CA/+^ Pten^Δ/+^ Sox2^fl/fl^*, *Braf^CA/+^ Pten^Δ/+^ Sox2^fl/+^*, *Braf^CA/+^ Pten^Δ/+^ Sox2^+/+^*mice were treated with tamoxifen at p2-4, a marked *Sox2* gene dosage-dependent decrease in tumor incidence was observed (Fig 3D, Fig S4D). Immunohistochemistry demonstrated that tamoxifen treatment successfully deleted *Sox2* expression in tumor cells (Fig S4E). These results indicate the path to tumor formation taken by *Braf^CA/+^ Pten ^Δ/+^* cells differs substantially from that taken by more heavily mutated melanocytes. Not only does the former type of tumor develop more slowly^13^ (Fig 1B) and display little pigment (Fig 1C), its development requires *Sox2* and thus the participation of a *Sox2*-expressing cell type. As LNM cells expressed the highest levels of *Sox2*, and are present even prior to tumor formation, these results raised the possibility that LNM cells play a role in tumor initiation.

### Comparative Analysis of Single-Cell RNA-seq Data between Mouse and Human Tumors

Unlike mouse models, in which multiple oncogenic mutations are usually introduced simultaneously, the order in which mutations (as well as other heritable changes) arise in human melanomas is unknown, and potentially differs from individual to individual. This prompted us to compare the cellular states that we detect in *Braf^CA/+^*and *Braf^CA/+^ Pten ^Δ/+^*mouse melanomas with those described for human melanoma samples, several of which have now been subjected to analysis by single cell RNA sequencing ^27^, as well as those identified in more heavily-mutated mouse melanomas. Although the direct comparison of gene expression states across species can be problematic, we took advantage of the fact that a variety of gene expression signatures have been proposed as markers of human melanoma cell states.^60^ To facilitate the mapping of agreement with these signatures onto our scRNAseq data, we developed a “membership score” pipeline that differentially weights the contributions of genes according to their degree of differential expression among the neural crest-derived cells in our samples (see Methods). Given a set of genes, and a collection of cells, the method assigns each cell a score representing an aggregate of how far above or below the average each gene’s expression is. To minimize batch artifacts, the neural crest-derived cells of each tumor genotype (Albino *Braf^CA/+^*, Albino *Braf^CA/+^ Pten^Δ/+^,* and black *Braf^CA/+^ Pten^Δ/+^*) were analyzed separately. Cells were displayed on individual UMAPs, and five clusters highlighted: principal tumor cells, proliferating tumor cells, melanocytes, Schwann cells, and LNM cells (Fig 4A, Fig S5A). Membership scores were calculated on a cell-by-cell basis for signatures (Table S10) that have been reported for treatment-naïve human melanoma biopsy^27^ (including patients with both primary and metastatic human melanomas), patient-derived xenografts grown in immune deficient mice and subjected to *Braf*-inhibitor treatment^31,60^, and other mouse melanoma models (*Nras^Q61K/+^ Ink4a^Δ/Δ^* and *Braf^CA/+^ Pten^Δ/Δ^*) ^27^.

**Figure 4.**
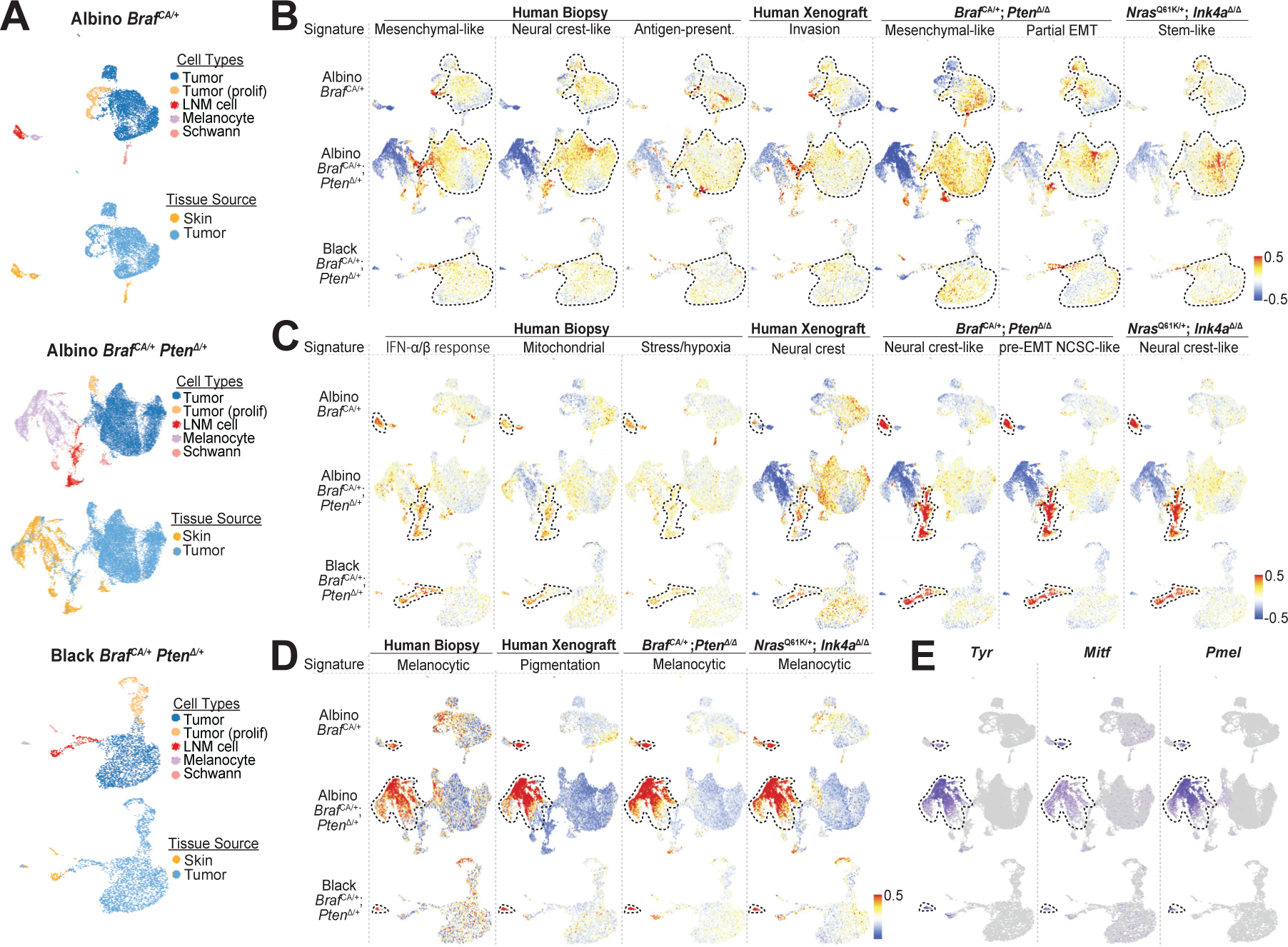
Signatures of neural crest-derived populations are conserved in mouse and human melanoma. **A.** scRNA-seq subclustering (top) and tissue source distribution (bottom) of neural crest-derived clusters from three genotypes: Albino *Braf*^CA/+^(21,600 cells), Albino *Braf*^CA/+^; *Pten*^Δ/+^ (5973 cells), and Black *Braf*^CA/+^; *Pten*^Δ/+^ (3926 cells). **B-D.** Every cell was assigned a score for how well its gene expression fit a published gene expression signature^27,31,60^, relative to the other neural crest-derived cell types (see Methods and Table S10) and these were overlayed on the UMAPs for each of the tumor genotypes (red=strong agreement; blue=poor agreement). Gene signatures that aligned primarily with principal tumor cells are shown in B; those that aligned strongly with LNM cells are shown in B and those that aligned with melanocytes are shown in C. Black dashes outline cell-types in the individual datasets. **E.** Feature plots of expression of selected pigmentation genes in the three tumor datasets. EMT: Epithelial-mesenchymal transitions. NCSC: Neural crest stem-cell.

Mapping these scores onto the UMAP plots in Figure 4 revealed similarities between the cell types identified in our mice and these signatures. Specifically, principal tumor cells had features resembling the “neural crest-like”, “antigen-presenting”, and “mesenchymal” signatures of human melanomas; the “invasion” signature of human xenografts; the “mesenchymal” and “partial-EMT” signatures of *Braf^CA/+^ Pten^Δ/Δ^* mouse tumor cells, and the “stem-like” signature of *Nras^Q61K/+^ Ink4a^Δ/Δ^* mouse tumor cells (Fig 4B). In contrast, LNM cells most closely resembled the “Interferon-alpha-beta response”, “mitochondrial”, and “stress” sets of human melanomas; and the “neural crest” and “neural crest stem cell like” (NCSC-like) signatures of xenografts and other mouse models (Fig 4C), signatures that overlapped only weakly with principal tumor cells.

Although both LNM and principal tumor cells expressed pigmentation genes at low levels, there was scant pigment visible on the surface of black *Braf^CA/+^ Pten^Δ/+^* tumors (Fig 1C), a phenomena described by others^62^, which motivated us to examine whether gene expression of any of the cells in these tumors aligned with the *Mitf*-high pigmentation signature that has been identified in other mouse and human melanomas. Membership scoring showed that *Mitf*-high “melanocytic” signatures identified by others in human biopsies, xenografts and pigmented mouse melanomas were observed mainly in skin melanocytes, only very weakly detected in principal tumor cells, and essentially absent from LNM cells in our tumors (Fig 4D-E).

### Prediction of Transient Cellular Dynamics Between LNM Cells and Principal Tumor Cells

Despite being distinct from principal tumor cells, several characteristics of LNM cells suggest they play a role in tumor formation and growth—for example, the importance of *Sox2*, which is most strongly expressed in LNM cells (Fig 3C), in tumor formation (Fig 3D); the persistence of LNM cells during serial transplantation (Fig 3A-C); and the expression within LNM cells of gene signatures associated with human and mouse melanoma cell states (Fig 4C). Two approaches were taken to test the possibility that LNM and principal tumor cells represent potentially interconverting states.

First, we applied RNA velocity analysis, using scVelo^63^, to the neural crest-derived cells of individual tumors (Fig 5B-D). RNA velocity uses levels of spliced and un-spliced transcripts to infer cell state dynamics, ordering cells based on the principle that high proportions of unspliced reads indicate recently induced genes while low proportions indicate genes recently turned off. We also analyzed cells using MuTrans, a pipeline that uses dynamical systems concepts to infer transient cells and cell-fate transitions from snap-shot single cell transcriptome datasets ^64^ (Fig 5E-H). To minimize batch-related artifacts, analyses were performed separately on individual black *Braf^CA/+^ Pten^Δ/+^* tumor samples. To increase the probability of detecting transitions involving LNM cells, we also analyzed two sets of tumors obtained after rounds of serial transplantation, as LNM cells are especially abundant in such samples (Fig 3B-C).

**Figure 5.**
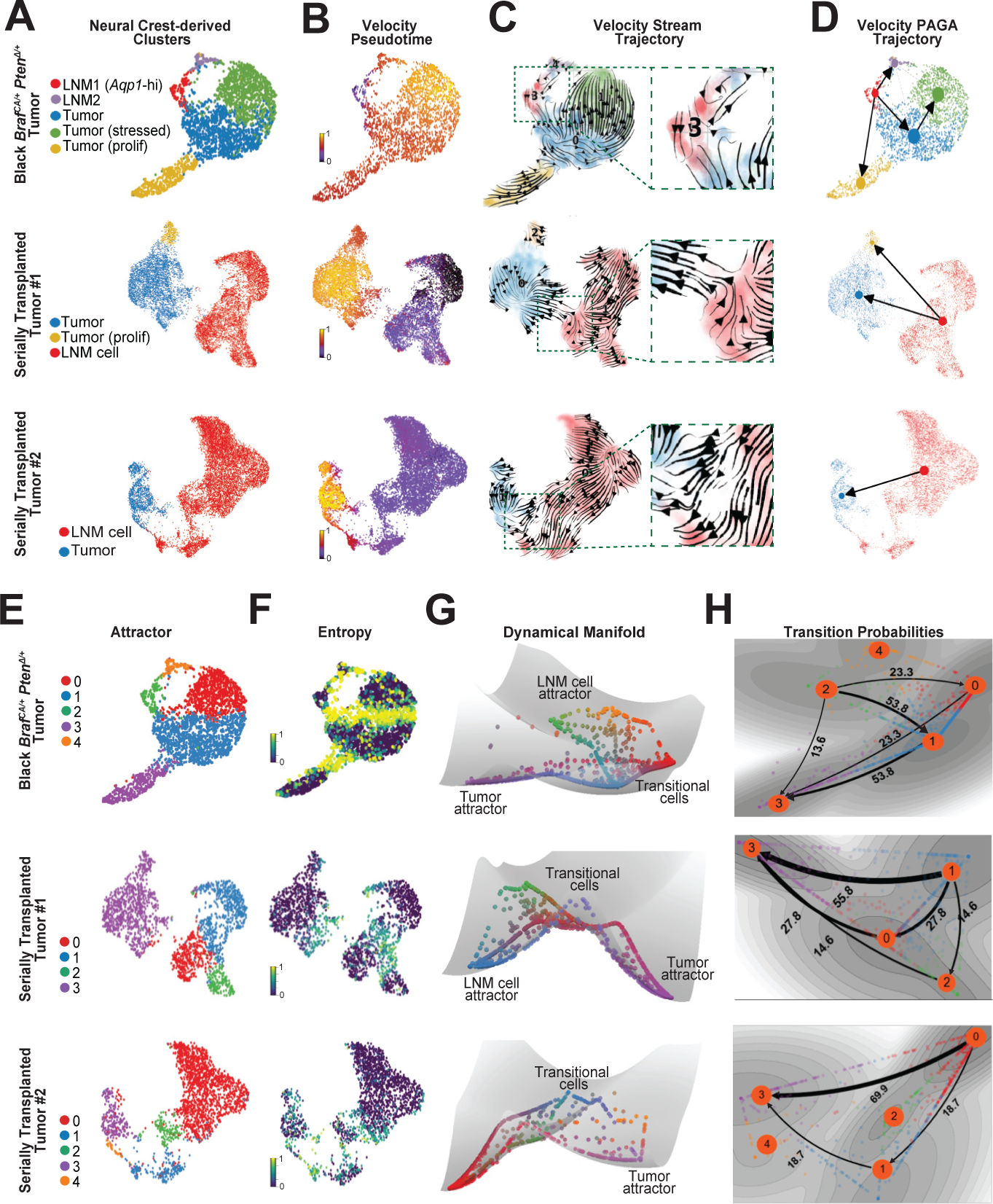
Evidence that LNM cells are transitional cells. **A.** As described in Fig 3A, tumors derived from Black *Braf*^CA/+^;*Pten*^Δ/+^ were serially passaged by transplantation into NSG (immune deficient) mice. Tumors from one parental Black *Braf*^CA/+^;*Pten*^Δ/+^ genotype mouse and two serially transplanted NSG mice (derived from a single tumor) were subjected to scRNA-seq, analyzed separately, and the neural crest-derived cells were sub-clustered accordingly (top panel: 2686 cells from the parental Black *Braf*^CA/+^;*Pten*^Δ/+^ tumor; middle and bottom panels: 10662 and 10965 cells from serially transplanted NSG tumors, respectively). **B-D.** RNA velocity analysis predicts fate decisions of individual cells in A using (B) Velocity pseudotime, (C) Velocity embedding stream trajectories (note the *Aqp1*-hi LNM-to-tumor transition in insets), and (D) Partition-based graph abstraction (PAGA), a topology clustering trajectory method with arrows summarizing directionality from *Aqp1*-hi LNM to tumor cells. **E.** Attractors (stable states) were identified by applying MuTrans to the same data sets. **F.** Larger entropy values suggest a more transient cell state. **G.** The dynamical manifold constructed by MuTrans, with potential wells representing stable attractors, and individual cells mapped onto the landscape **H.** Transition path analysis calculates predicted relative rates of transition between stable attractor states. Note the transition path from the LNM cell attractor to Tumor attractor. The numbers along paths indicate the proportion of total transition flux, with larger values suggesting higher likelihood of transition.

In every sample both RNA velocity and MuTrans independently supported the conclusion that transitions occur between LNM and principal tumor cells (Fig 5B-D, Fig 5H). In primary tumors, such transitions appeared to involve a subset of LNM cells expressing high levels of *Aqp1* (identified in Fig 5A as “LNM1”). Among all possible transitions identified by MuTrans, the path from LNM to primary tumor was consistently identified as the most likely of all possible transitions (Fig 5B-D, Fig 5H).

## DISCUSSION

Genetic heterogeneity, epigenetic modifications, and a panoply of cell states are characteristic of melanoma^32^, yet the background rate of mutation in human skin^65,66^ makes it difficult to ascertain relationships between cell state landscapes, individual mutations and non-genetic factors. In addition, melanomas in individual patients can differ markedly in histological characteristics, and can include the absence of pigmentation^67^ as well as the expression of different degrees of “neural” phenotype^68^.

Mouse models have yielded important insights into how melanomas arise^22^. In mice, combining gain of function oncogene mutations (*Braf* and *Nras*) with complete loss of function mutations in tumor suppressor genes (*Cdkn2a, Pten, Ink4a*) provides an efficient route to the rapid generation of melanoma^69,70^, but by the same token may not shed light on diversity of routes by which melanocytes transition to malignancy. Here we concentrated on melanomas generated either solely by *Braf* activation, or by *Braf* activation in combination with heterozygous loss of function of *Pten*. Such melanomas arose slowly and rarely, with kinetics indicating the requirement for a low-probability random event. Interestingly, we found no evidence that a de novo mutation (including loss of heterozygosity for *Pten*) accounted for that event, suggesting the need for an epigenetic transition. The tumors produced grew slowly and were hypomelanotic, unlike the rapidly-growing, pigmented tumors that arise when greater numbers of mutations are combined. Despite this, signatures of cell subsets found in the melanomas described here overlap with those described in subsets of cells in more heavily mutated mouse models. In particular, the LNM cells described here overlapped substantially in gene expression with very lowly-pigmented “neural crest” and “neural crest stem cell-like” signatures seen in other models. In human xenograft models, this slow-growing state is seen in substantial numbers only after treatment with *Braf* inhibitors, which eliminates the major, highly-pigmented tumor cells of these xenografts. Intriguingly, it has been proposed that these cells serve as an intermediate on the way to a final, *Braf*-inhibitor resistant tumor cell type^31,61^. Based on comparisons with other studies, we propose a model in which two pathways exist for melanomagenesis (Fig 6).

**Figure 6.**
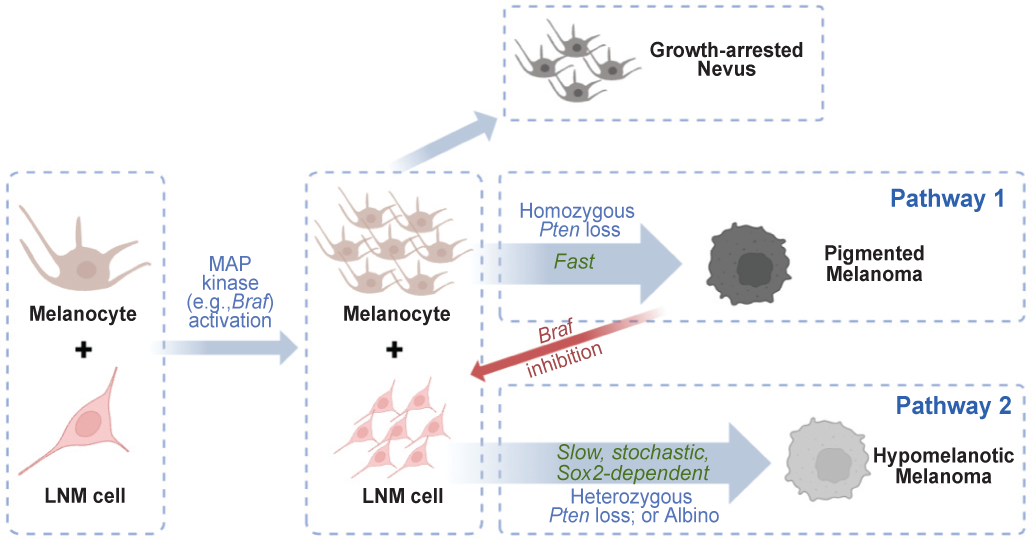
Proposed pathways to melanomagenesis. Skin contains both pigmented melanocytes and rare, poorly-pigmented LNM cells. Upon *Braf* or *Nras* activation, both populations expand, but melanocytes arrest as nevi, unless loss-of-function mutations in tumor suppressor genes allow them to develop rapidly into pigmented melanomas (Pathway 1). Otherwise, *Braf*-activated LNM cells can undergo rare, stochastic transitions to produce scantly pigmented melanomas (Pathway 2). The probability of such transitions depends on host pigmentation status, *Sox2* expression, and whether a single allele of *Pten* has become inactivated. LNM cells persist in both types of tumor, and in response to *Raf/ Mek* inhibition, they are thought to provide a source for the emergence of drug-resistant cells^31^.

We begin with the assumption that skin contains both pigment-producing melanocytes and poorly pigmented LNM cells—the latter potentially corresponding to various types of neural crest or melanocytic precursor cells that have been described in normal skin ^40,71^. Under normal conditions, these cell types may exhibit a precursor-product relationship, or even interconvert reversibly, but this hypothesis remains to be tested. Introduction of an activating *Braf* or *Nras* mutation into melanocytes and/or LNM cells is sufficient to drive the rapid formation of growth-arrested nevi. In mice at least, *Braf* activation also leads to an expansion of LNM cells (Fig S2C). If there is also a sufficient loss of tumor suppressor gene function—in human melanoma this could reflect prior mutation accumulation, whereas in mice this requires engineered gene inactivation—then rapid progression to a pigmented tumor occurs.

In contrast, if no additional mutations are present, a rare, stochastic event allows for the transition of *Braf* transformed LNM cells into slow-growing, hypomelanotic tumors. Although the molecular nature of that event is as yet unknown, it can be accelerated either by an albino genotype, or the loss of a single allele of *Pten*, suggesting it is sensitive to even subtle changes in the growth properties of LNM cells. It is possible that the stemness-associated transcription factor *Sox2,* which is required for tumor formation by this route (Fig 3), itself plays a role in the transition to malignancy, however it is also possible that *Sox2* is simply required for LNM cell growth or survival.

Although two independent paths to melanomagenesis are presented in Figure 6, there are good reasons to believe both may operate together in human tumors. In particular, it was reported that, when human xenografts were subjected to Braf inhibition, a distinct subset of cells survived and acted as a precursor to the development of inhibitor-resistant tumors. This subset was described as slow growing, poorly-pigmented, and characterized by a neural crest-like signature closely resembling the gene expression pattern of LNM cells (for example, such cells are marked by *Aqp1*). Although they were described as arising in response to inhibitor treatment, it seems likely these cells pre-existed as a minor population; indeed, a minor population with signatures overlapping with LNM cells can be found in primary human melanomas (Fig 4C), as can *Aqp1*+ cells histologically^72^. Whether this population grows alongside pigmented tumor cells, or is actively generated by them, remains to be determined. Interestingly, a distinctive marker of LNM cells is the expression of *Igf1* (Fig 2D, S2B), which encodes a growth factor that stimulates melanoma cell growth ^73^, raising the possibility that LNM cells may also have a trophic role in sustaining tumors.

The nature of the stochastic event required for producing hypomelanotic melanoma remains to be determined. One caveat to our attempt to rule out a mutational cause is that whole genome sequencing will not necessarily detect all structural variants, nor could we eliminate the possibility that a different mutational route was followed in each tumor. Still, it seems most likely that the explanation is epigenetic, similar to the slow state switching that is thought to play a role in melanoma progression and persistence^74^. To our knowledge, this is the first suggestion that such switching might also play a role in melanoma initiation.

The observation in the present study that a change in pigment background markedly influenced the likelihood of tumor development (Fig 1A-B) is consistent with both experimental literature and epidemiologic data on the incidence of melanoma in light skinned versus dark skinned individuals ^37^. In mice, pheomelanin-producing melanocytes apparently have a higher transformation potential than eumelanin producing melanocytes^62^, and eumelanin can inhibit melanomagenesis^75^. Further, the absence of melanin, despite having little effect on the mutation rate in melanocytes, increases the aggressiveness of tumors in *Nras* melanoma models ^76^.

Melanoma can be cured if detected early and removed^77^, but it is often difficult to distinguish from benign lesions using clinical examination, histology, ^78^ or DNA alterations ^6,79^. It is intriguing to note that, in the present study, LNM cells accumulated in *Braf^CA/+^ Pten^fl/+^* and *Braf^CA/+^* mice even in skin in which tumors had not yet developed (or could not do so). To the extent these cells play a necessary role in the development of melanoma by at least one pathway, and may also be involved in the development of resistance in melanomas generated by the other pathway^31^ (see above), their detection in biopsies may represent a clinically useful marker of malignant potential. Furthermore, to the extent that these cells can ultimately act as a progenitor to tumor cells, they may also represent a target for development of preventive therapies. Indeed, the fact that malignant transformation of these cells appears to involve an epigenetic, rather than genetic, transition raises the possibility that a wider range of therapeutic modalities could be explored than are usually considered in skin cancer prevention, which is currently focused on reducing mutation burden.

## MATERIALS AND METHODS

### Key resources table

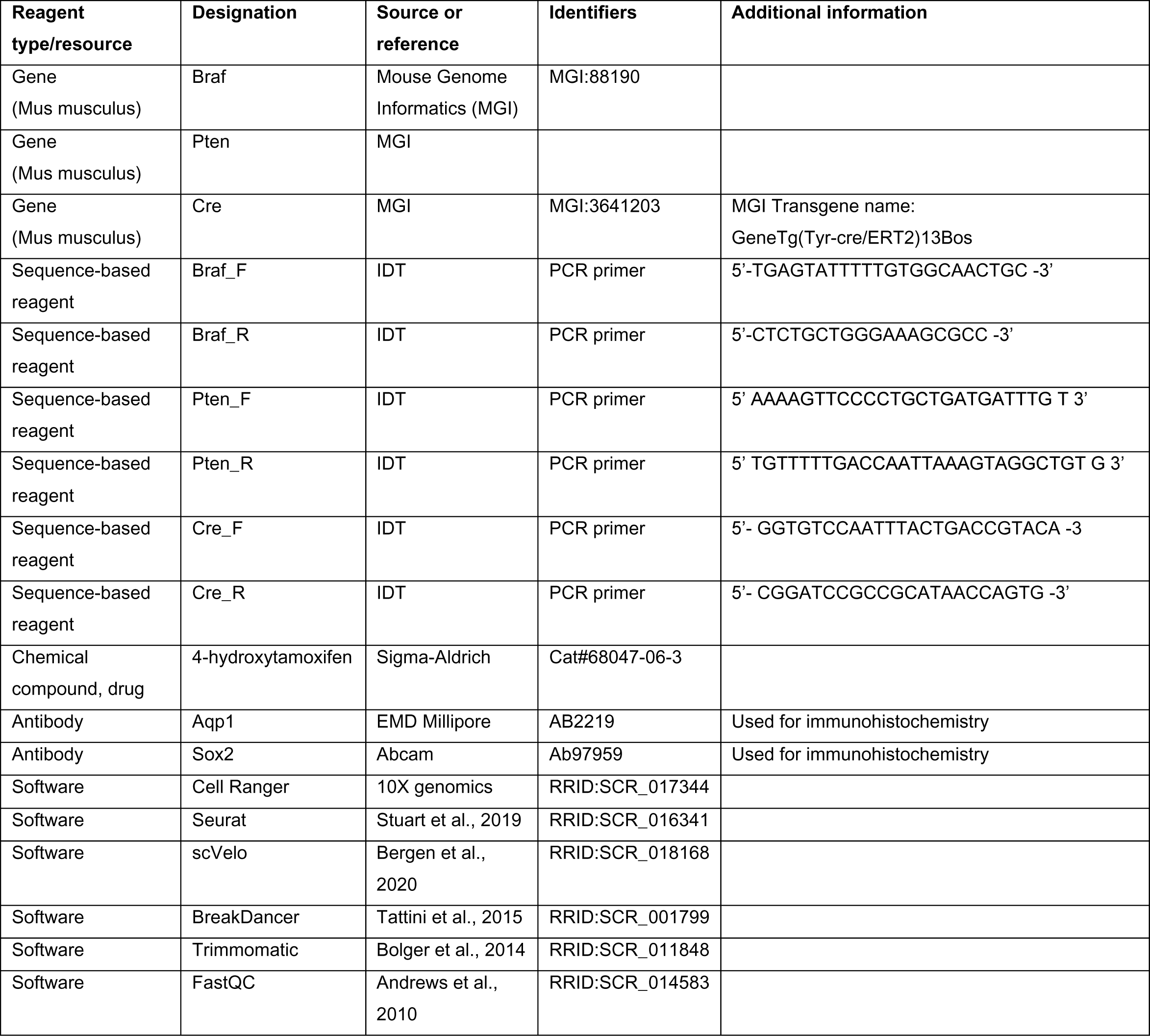

### Generation of *Braf*^CA/+^ melanoma and nevi bearing mice

*Braf*^CA-fl/+^, Tyr-CreER (C56BL/6) mice (RRID:MGI:5902125) were genotyped by PCR as previously described (Bosenberg et al., 2006; Dankort et al., 2007). The primers used in this study are: *Braf* forward 5’-TGAGTATTTTTGTGGCAACTGC -3’, *Braf* reverse 5’-CTCTGCTGGGAAAGCGCC -3’, *Pten* forward 5’-AAAAGTTCCCCTGCTGATGATTTGT -3’, *Pten* reverse 5’-TGTTTTTGACCAATTAAAGTAGGCTGTG -3’. *Cre* forward 5’-GGTGTCCAATTTACTGACCGTACA -3’ and *Cre* reverse 5’-CGGATCCGCCGCATAACCAGTG -3’. Topical 4-hydroxytamoxifen (4-OHT; 25 mg/mL or 75 mg/mL in DMSO; 98% Z-isomer, Sigma-Aldrich) was administered to pups on their back at ages P2-P4. Mice were euthanized if the volume of their tumors exceeded 10% of total body volume, if tumors were significantly ulcerated, if mice were moribund, if they lost weight, if they were lethargic, or if they were unable to ambulate. All mouse procedures were approved by UCI’s IACUC.

### Tissue Harvest for Whole-genome Sequencing

Melanoma-bearing albino *Braf*^CA/+^ mice (n=3, aged postnatal day 90 to 325) were euthanized, shaved, and depilated. Tumor (25mg), skin (25mg), and spleen (10mg) were then collected from each mouse and cut into small pieces with a scalpel, and the fat scraped off from the underside of the skin. Collected tissues were then stored in -80°C freezer until sample preparation day. DNA materials were extracted using the Qiagen blood and tissue Kit, following protocols described for purification of total DNA from animal tissues. Briefly, approximately 25 mg of tissue was obtained and digested with ATL buffer and proteinase K. Ethanol was added to the lysate and passed through a DNeasy spin column. DNA was eluted with AE buffer.

### Library Preparation for Whole-genome Sequencing

Library construction was performed using the NEXTflex Rapid DNA-Seq kit v2 and the NEXTflex Illumina DNA barcodes. Using the Covaris S220, 50ng of gDNA was sheared using settings to target 400bp. The sheared gDNA was end repaired and adenylated. The reaction mixture was cleaned up using AMPure XP magnetic beads and Illumina barcoded adapters were ligated onto the blunt-end/adenylated product. The adapter ligated product was cleaned using AMPure XP beads and then amplified for adapter ligated products using 5 cycles of PCR. The resulting library was cleaned with AMPure XP beads with double sided size selection and then quantified by qPCR with Kapa Sybr Fast universal for Illumina Genome Analyzer kit. The library size was determined by analysis using the Bioanalyzer 2100 DNA HighSensitivity Chip. The library was sequenced on the NovaSeq 6000 Sequencer, using an S4 flowcell chemistry and PE100 cycles with additional cycles for the index read. Sequences were obtained as paired-end 100 bp reads. The NovaSeq control software was v1.6.0 and the real time analysis software (RTA v3.4.4) converted the images into intensities and base calls. Post-processing of the run to generate the FASTQ files was performed at the UCI Institute for Genomics and Bioinformatics (UCI IGB).

### Whole Genome Sequencing bioinformatic analysis

<u>SNV analysis</u>, Raw reads were transferred, and quality analyzed using *FastQC* tool (v0.11.9). Low quality bases and adapter sequences were trimmed using *Trimmomatic* (v0.39)^80^. Trimmed reads were then aligned to the mouse reference genome (build mm10) using the Burrows-Wheeler Aligner (*BWA mem*, v0.7.12)^81^. Duplicated reads were removed using *Picard tools MarkDuplicates* (v1.130). Local realignment and base quality recalibration was done on each chromosome using the Genome Analysis Toolkit (*GATK* v4.0.4.0) best practices. SNVs and indels were detected and filtered using GATK with HaplotypeCaller function. The output files were generated in the universal variant call format (VCF).

Somatic mutations were identified and filtered using mutect2(v4.1.3.0)^82^ and bcftools (v1.15.1)^83^. To estimate the fraction of DNA in each sample that was derived from *Braf^CA/+^*-expressing cells (as opposed to fibroblasts, immune cells, etc.), we identified all reads that mapped to the loxP sequence and that included at least three bases on either side. From the sequences up- and downstream of loxP we were able to uniquely assign each read to one of three categories: an un-recombined 5’ loxP site, and unrecombined 3’ loxP site, and a recombined loxP site. Unrecombined cells should contain one each of the sequences of the first two types, whereas recombined cell should contain a single sequence of the third type. Thus, the ratio of the observed number of sequences of the third type to the average of the number of sequences of the first two types provided an estimate of the ratio of recombined to unrecombined cells.

<u>CNV analysis</u>, BreakDancer (v1.1.2)^84^ was used to analyze mapping results from whole genome sequencing and classify structural variations in five categories: deletion, insertion, inversion, intrachromosomal translocation and interchromosomal translocation. Genomic regions with statistically significant amount of anomalous read pairs (split reads, depth) are considered supporting structural variations breakpoints. A Poisson model is implemented to incorporate the number of supporting anomalous reads, the size of the region and the genomic coverage to compute a confidence score. The default set of parameters was used for BreakDancer.

We followed the recommended BreakDancer protocols^85^ and analyzed the matched tumor, skin and spleen samples for each of the 3 animals. BreakDancer was run 3 times, each with 3 genomes from the same animal. In order to get somatic CNVs, the identified breakpoints were then filtered so that only those from the tumor are preserved (Tumor-skin-spleen). The 3 identified CNV lists from the 3 animals were then annotated with gene symbols if overlapping with the gene body. The 3 resulting gene lists were intersected to get plausible recurrent somatic CNVs.

### Cell isolation for single-cell RNA sequencing

Nevi-bearing *Braf*^WT^, *Braf*^CA/+^, and *Braf*^CA/+^ *Pten* ^Δ/+^ mice were harvested at P30 and P50 (n = 3 of each genotype). Melanoma bearing *Braf*^CA/+^ (albino coat color), *Braf*^CA/+^ *Pten* ^Δ/+^ (albino and black coat color), and transplanted tumors were harvested when tumors reached a size that affected the health of the animal, as described above, complying with IACUC regulations. The mice were euthanized, shaved, and depilated. For non-tumor bearing samples, a 2 by 3 cm section of the back skin was retrieved, and the fat scraped off from the underside. For melanoma-bearing samples, tumor tissue, the skin sample on top of the tumor piece (termed “tumor skin”) were collected, with the mouse-matched adjacent skin next to the tumor (termed “tumor-adjacent skin”) as sample-matched control.

The tumor or skin sample was then physically minced, and suspended in a gentle-macs C-tube dissociation buffer (5mL of RPMI, 50uL of liberase 0.25 mg/mL, 116uL of Hepes 23.2 mM, 116uL of Sodium Pyruvate 2.32 mM, 500uL of Collagenase:Dispase 1 mg/mL) for 50 min at 37°C incubation with gentle agitation at a speed of 85-90rpm. After initial 50 minute digestion i, 10uL of DNaseI (23.2uL) was added for another 10 min of 37°C agitated incubation and then inactivated with 400uL of fetal bovine serum and 10uL of EDTA (1 mM). The tissue suspension was further dissociated mechanically with GentleMACS by running the setting m_imptumor_04.01 twice. The digested suspensions were filtered twice through a 70 mm strainer and dead cells removed by centrifugation at 300 x g for 15 min. The live cells were washed with 0.04% UltraPure BSA:PBS buffer, gently re-suspended in the same buffer, and counted using trypan blue and cell counter.

### Library preparation for single-cell RNA sequencing

Libraries were prepared using the Chromium Single Cell 3’ v2 protocol (10X Genomics). Briefly, individual cells and gel beads were encapsulated in oil droplets where cells were lysed and mRNA was reverse transcribed to 10X barcoded cDNA. Adapters were ligated to the cDNA followed by the addition of the sample index. Prepared libraries were sequenced using paired end 100 cycles chemistry for the Illumina HiSeq 4000. FASTQ files were generated from Illumina’s binary base call raw output with Cell Ranger’s (v2.1.0; RRID:SCR_017344) ‘cellranger mkfastq’ command and the count matrix for each sample was produced with ‘cellranger count‘.

### Single-cell RNA sequencing analysis

47 samples *from the genotypes: wild type; Braf^CA/+^; Braf^CA/+^ Pten^Δ/+^; and Pten^Δ/+^* were aggregated using the Seurat built-in Merge function to produce one count matrix. Downstream bioinformatic analysis was conducted using Seurat^86^. For quality control, cells with fewer than 200 detected genes and genes detected in less than 3 cells were discarded. We calculated the percent mitochondrial gene expression and kept cells with less than 15% mitochondrial gene expression, and cells with fewer than 4000 genes per cell. A total of 345,427 cells from the whole skin samples passed the quality control. Each cell was then normalized using scTransform. In the final preprocessing step, we regressed out cell-cell variation driven by mitochondrial gene expression and the number of detected UMI. To identify cell-type clusters, principal component analysis using highly variable genes, Louvain clustering, and visualization with Uniform Manifold Approximation and Projection (UMAP) was used. The Euclidean distances between clusters was calculated using the average principal component (PC) embedding for each cluster, from the top 10 PCs.

### Histopathology and Immunostaining

For immunohistochemistry, formalin fixed paraffin embedded sections were sectioned around 5mm - 8 mm thick onto poly-L-lysine coated slides, under tissue water bath temperature 31 to 37°C. Slides were deparaffinized with Xylene, and dehydrated in a series of increasing concentration of ethanol washes. Antigen retrieval was performed with 10 mM citric acid buffer at pH 6.0 for 50 min in a 80°C water bath, then left to cool overnight at room temperature. All samples were next incubated with first antibody overnight at 4°C. Samples were washed and incubated with the appropriate secondary antibody. H and E staining was performed using standard histopathological methods^87^.

### Congenic Tumor Transplantation

Melanoma-bearing *Braf*^CA/+^ *Pten* ^Δ/+^ donor mice were euthanized, shaved, and depilated when tumor reached around 10% of body weight. After tumor was collected, it was cut into pieces (weighing around 0.02g-0.06g) in a cake-slice pattern to preserve tumor structural heterogeneity; this was performed in a biosafety cabinet to ensure a sterile environment. The pieces were then washed twice with 70% EtOH, and once with 1x PBS, each for 10 seconds. They were then transferred to RPMI + 10% FBS media on ice until transplantation.

NSG recipient mice were anesthetized using Isoflurane. The tumor piece was transplanted to the back flank skin, by cutting a small incision and slipping the tumor tissue in together with 100uL of Matrigel, after the mouse was shaved and sanitized with 100% EtOH and chlorhexidine diacetate disinfectant solution. Transplantation shams were also included (n=1-2 per group) by only transplanting 100uL of Matrigel to the mouse back. 10uL of lidocaine was applied to the wound for consecutively three days after the procedure. Tumor size was measure weekly.

### Generation of *Braf*^CA/+^ *Pten^Δ/+^ Sox2*-deficient mice

*Sox2*-floxed mice were purchased from Jackson laboratories (STOCK Sox2tm1.1Lan/J strain#:013093) and crossed to *Braf^CA-fl/+^ Pten^fl/+^* mice that had been backcrossed into a pure C57Bl6 background for greater than six generations to generate *Braf^CA-fl/+^ Pten^fl/+^ Sox2*^fl/fl^ mice. Induction of tumors in these mice were performed as described above.

### Membership score analysis

To quantify how well cell types identified by single cell RNA sequencing fit a proposed gene expression signature, we first converted gene expression values to modified, corrected Pearson residuals, as described^88^, using empirically determined estimates for the coefficient of variation of biological gene expression noise of 0.55, 0.65, 0.65 respectively, for *Albino Braf^CA/+^*, *Albino Braf^CA/+^ Pten^Δ/+^*, and *Black Braf^CA/+^ Pten^Δ/+^* samples, respectively. Residuals were then standardized, essentially converting them to z-scores. The average of the z-scores for all of the genes in any proposed gene signature was then calculated for each cell, and this value was plotted as a color-score on the UMAP diagram of the cells.

### RNA velocity analysis

The scVelo package^63^ (version 0.2.5) was used for RNA velocity analysis. To estimate the velocity based on unspliced and spliced counts, we applied “dynamical” mode which uses differential equations to model the transient dynamics of gene expression and splicing, with other parameters set as default. Pseudotime was calculated by the “velocity_pseudotime” function with default parameters. The streamlines of velocity were visualized using “velocity_embedding_stream” function with parameter density set as 1. Cell lineage analysis was then performed by Partition-based graph abstraction (PAGA) (provided in scVelo package) based on the calculated velocities.

### MuTrans analysis

For MuTrans analysis^64^, we used all cells in the Black *Braf^CA/+^ Pten*^Δ/+^ sample and also subsampled 20% cells from the two transplanted NSG mouse tumor samples. The number of attractors were determined by EPI (eigen-peak index) of MuTrans. The cellular random walk was constructed using cosine similarity of gene expression as the input to MuTrans. The dynamical manifold was constructed using default parameters based on UMAP dimension reduction coordinates as the initial values. Transition gene analysis was performed using the “GeneAnalysis” module in MuTrans, with parameter “thresh_de_pvalues” set as 5e^-4^ and all other parameters kept as default.

### Statistical analysis

Kaplan Meier survival curves were generated, and significance assessed using the log-rank test, an unpaired t-test, or a two-way ANOVA test. Other data were analyzed by GraphPad Prism6 statistical analysis software using an unpaired t-test or two-way ANOVA test. Significance levels were as indicated.

## Author contributions

Conceptualization: ADL, AKG

Data curation: HX, JS, CC, SST, RRV

Formal analysis: HX, JS, CC, JW, PZ, SST, RRV, ADL

Investigation: HX, JS, CC

Writing - original draft: HX, ADL, AKG

Writing - review and editing: HX, JW, PZ, ADL, AKG

Funding acquisition: ADL, AKG

Methodology:ADL, PZ, JW, QN

Project administration: ADL, AKG

## Acknowledgement and Funding

This work was supported by NIH CA217378, CA244571, and P30AR075047. HX was supported by CIRM under Award EDUC4-12822. RRV was supported by the UC Presidents fellowship and a FORD Foundation Fellowship. QN was partly supported by supported by NSF grants DMS1763272 and a Simons Foundation grant (594598). This work was made possible, in part, through access to the Genomics High Throughput Facility Shared Resource of the Cancer Center Support Grant (CA-62203) at the University of California, Irvine and NIH shared instrumentation grants 1S10RR025496-01, 1S10OD010794-01, and 1S10OD021718-01. Biorender was used for schematics.

