## supplemental figures for "Uncovering Minimal Pathways in Melanoma Initiation"

### Suppl. Figure 1

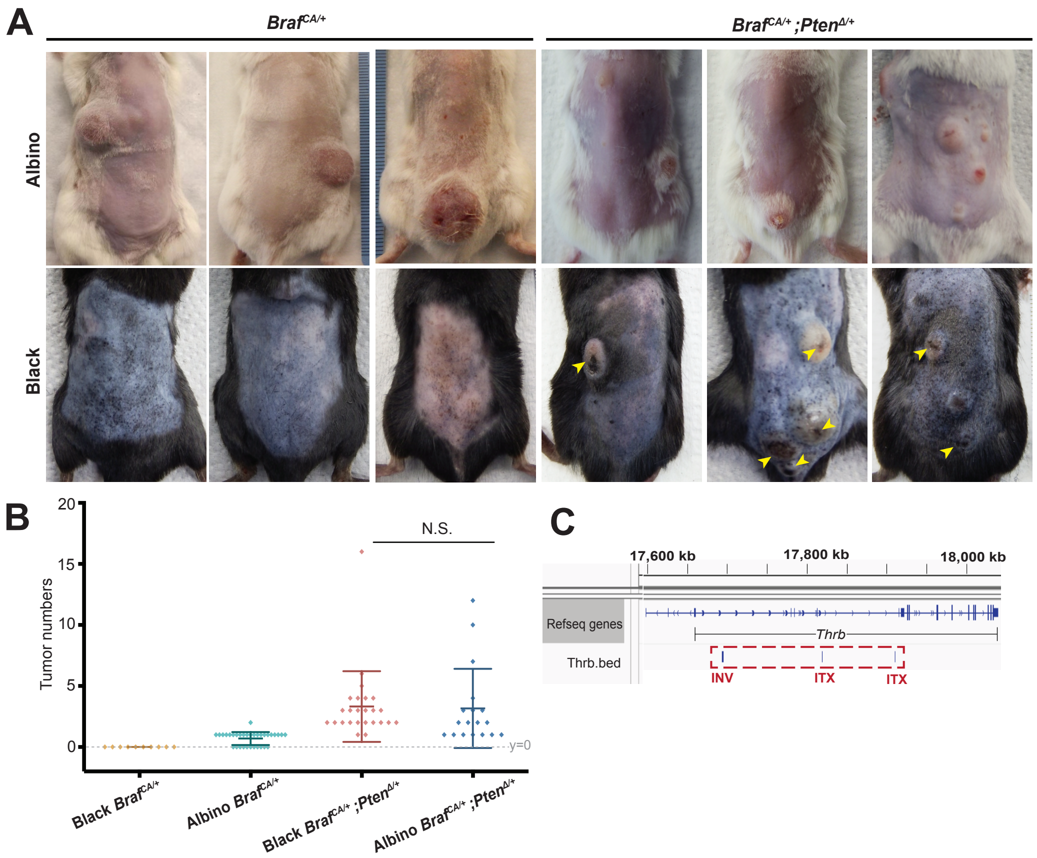


**Supplementary Figure 1. A-B.** Additional representative images (A) and numbers of tumors per animal (B) for *Braf*$^{CA/+}$ and *Braf*$^{CA/+}$; *Pten*$^{\Delta/+}$ models in different coat-color backgrounds. Arrows in panel A highlight the scant pigment observed associated with tumors in black mice. **C.** Mapping of *Thrb* structural variants described in Fig. 1D. Note the variant types are inconsistent and the variant regions do not overlap. INV: Inversion. ITX: Intrachromosomal translocation. *Thrb* expression in the tumor cells of these samples is only barely detectable, and is much lower than in other cell types.

### Suppl. Figure 2

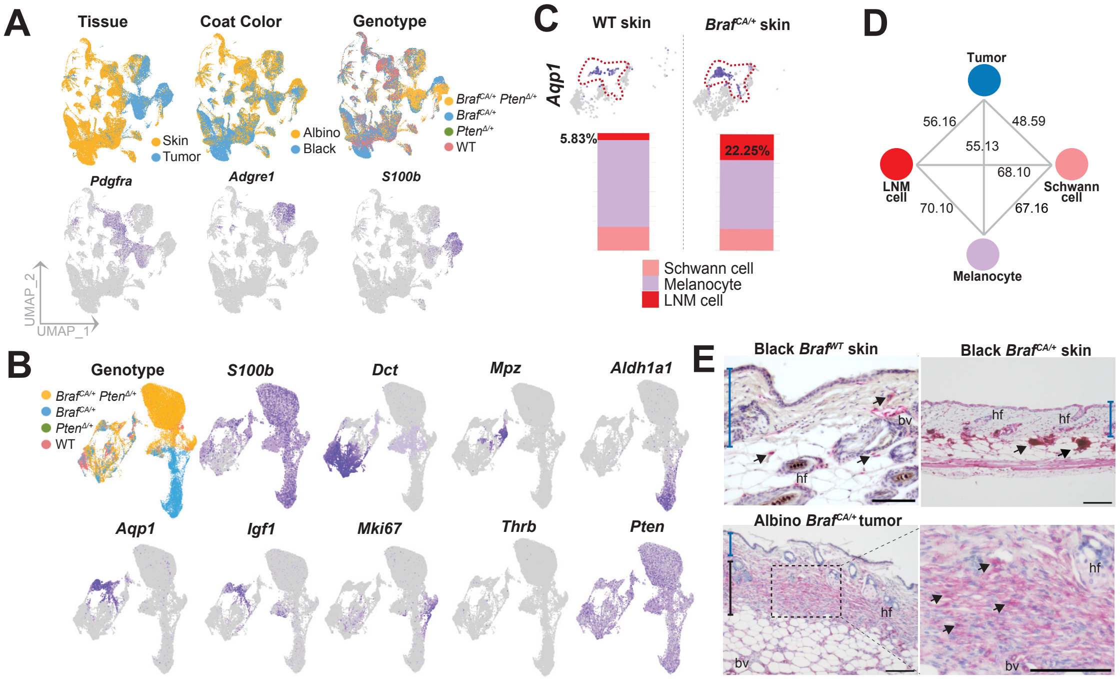


**Supplementary Figure 2. A.** Additional UMAP analysis of gene expression in the whole-skin single-cell RNA-sequencing of 345,427 cells shown in Fig 2B. Top: sample distribution labeled by tissue, coat color, and genotype. Bottom: Feature plots highlighting cell-types of interest based on known marker genes: Fibroblasts (*Pdgfra*), Macrophages (*Adgre1*) and Principal tumor cells (*S100b*). **B.** Additional visualization by genotype, and feature plots of the merged neural-crest-derived clusters (35,527 cells from Fig 2C), highlighting distinct cell-types. **C.** scRNA-seq feature plots of the LNM marker *Aqp1* and cell proportion plots in melanocyte cells across WT and *Braf*-mutant skin. Note the expansion of LNM cells in skin from black *Braf*$^{CA/+}$ mice (as tumors do not form in these mice, such cells cannot be tumor-derived) **D.** Euclidean distances calculated between NC-derived clusters using embeddings from the top 10 principal components (PCs). **E.** Immunohistochemistry of LNM cells (pink staining of *Aqp1+*; black arrows). Samples were counterstained with hematoxylin. Note the strong staining in the nevi of skin samples. Note that *Aqp1* straining spares the superficial dermis (blue bracket) but extends to the subcutis (black bracket) in tumors. hf: hair follicles. bv: blood vessels. Scale bar: 100um.

Suppl. Figure 3

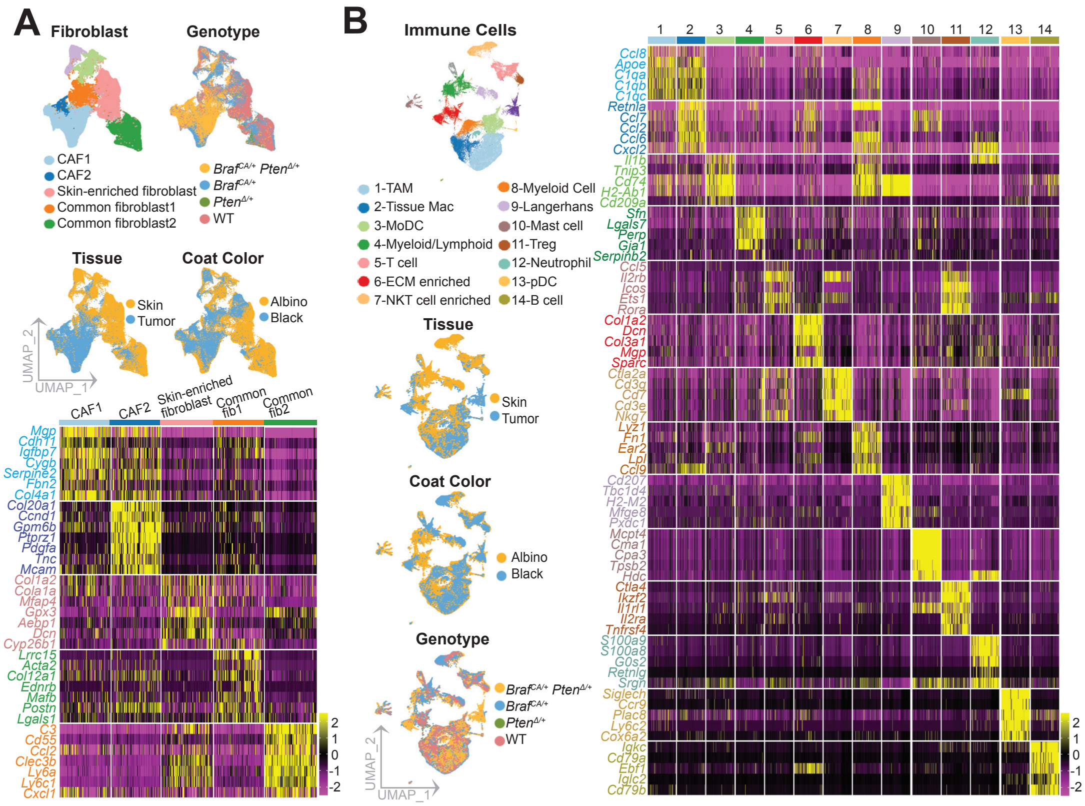


**Supplementary Figure 3.A-B.** Gene expression profiles of (A) 86,536 fibroblasts and (B) 51,747 immune cells associated with the skin and tumor microenvironments, as described in Fig 2B. Cells were subclustered and visualized by UMAP, and also labeled by tissue of origin, coat color, and genotype. CAF: cancer-associated fibroblast. TAM: tumor-associated macrophage. MoDC: Monocyte-derived dendritic cell. ECM: Extracellular matrix. Treg: Regulatory T cell. pDC: Plasmacytoid dendritic cell.

### Suppl. Figure 4

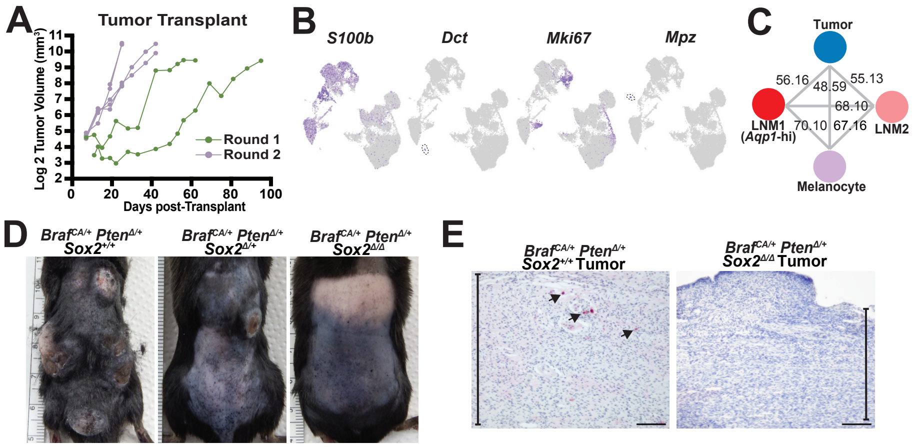


**Supplementary Figure 4. A.** Tumors were transplanted from one Black *Braf*$^{CA/+}$; *Pten*$^{\Delta/+}$ tumor into two NSG hosts in the first round and grown; and one primary transplanted tumor was harvested and serially re-transplanted into a second round of five NSG hosts. Growth curve for each tumor is recorded. Note that the time delay for tumor initiation is shortened in round 2. **B.** Feature plots highlighting neural crest-derived cell types: Principal tumor cells (*S100b*), melanocytes (*Dct*), highly proliferating cells (*Mki67*), and Schwann cells (*Mpz*). **C.** Euclidean distances calculated between NC-derived cell clusters in Fig 3B using embeddings from the top 10 principal components (PCs). **D.** Images from *Braf*$^{CA/+}$; *Pten*$^{\Delta/+}$ *Sox2*-depletion experiment. Note the absence of tumor when *Sox2* is eliminated by Tyr::Cre-ERT2. **E.** Immunohistochemistry of *Sox2* (pink staining), showing *Sox2* deletion in the sole *Braf*$^{CA/+}$; *Pten*$^{\Delta/+}$ *Sox2*$^{\Delta/\Delta}$ mouse tumor, compared to the positive staining (indicated by arrows) in a *Sox2*-wildtype tumor. Seconds are counter-stained with hematoxylin. Scale bar: 100um.

### Suppl. Figure 5

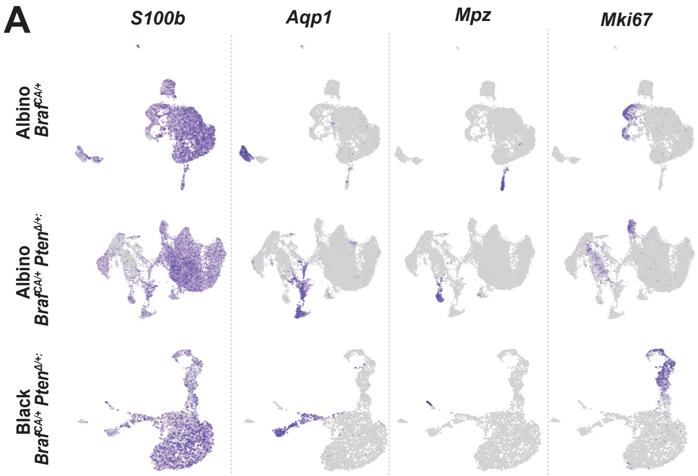


**Supplementary Figure 5.** A. ScRNA-seq feature plots of neural crest-derived clusters from three datasets in Fig 3A, highlighting distinct cell-types: Principal tumor cells (*S100b*), LNM cells (*Aqp1*), Schwann cells (*Mpz*), and a highly proliferative subset of principal tumor cells (*Mki67*).
